# Agro-morphological and phenotypic variability of sweet basil (*Ocimum basilicum* L.) genotypes for breeding purposes

**DOI:** 10.1101/2020.05.19.103861

**Authors:** Gulsum Yaldiz, Mahmut Camlica

## Abstract

The genus Ocimum is very complicated due to the presence of huge morphological variability along with genetic diversity. Basil (Ocimum basilicum L.) has pharmacological properties like headaches, coughs, diarrhea, constipation, warts, worms, kidney malfunction, and its potential as a therapeutic agent in treating various age-related diseases. The present investigation comprised of sixty-one genotypes of basil was undertaken to characterize the genotypes based on morphological and phenological features, herbs and essential oil yield of genotypes. A wide range of variations for traits like days to first cutting (56.92-101.6), plant height (13.67-71.90 cm), branch number (3.28-19.43 number/plant), fresh herb yields (12.94-274.11 g/plant), and essential oil yield (0.04-1.71%) were observed and can be useful for breeding purposes. PI 652070 and PI 296391 genotypes were found superior in case of the highest herbs yield as compared with other genotypes. Overall, in PI 358469 and Ames 32309 genotypes exhibited the highest essential oil content. The constellation analysis was conducted to investigate the genetic diversity of basil genotypes. According to the constellation plot analysis, leaf shape and color were evaluated in 2017 and most of the basil genotypes located in the same main group. In 2018, moonlight and dino cultivars located in the same cluster 1 (C1) with PI 141198 (US/Maryland) genotype and Georgia genotypes located in the same main group and they also took place in the sub-main group except Ames 32314 genotype depending on UPOV criteria. Each two years, Bolu genotype and midnight were found in the same main group.

## 1. Introduction

The genus Ocimum comprises more than 150 species that were reported throughout the world, but commercially important basil cultivars mostly belong to the species *O. basilicum*. Sweet basil (*Ocimum basilicum* L. x=n = 12) is an important essential oil crop in the Labiatae family, is an annual plant and is widely used as in food, pharmaceuticals, and cosmetics [1]. Also, the essential oil of obtained from its aerial parts contain linalool, eugenol, methyl chavicol, methyl cinnamate, ferulate, methyl eugenol, triterpenoids, and steroidal glycosides, which has high economic value because it contains important components, such as eugenol, chavicol, and their derivates [2–4]. Traditionally, basil has been used as a medicinal plant in the treatment of headaches, coughs, diarrhea, constipation, warts, worms, kidney malfunction [5], to cure malarial fever [6,7], as well as against mosquito vectors and plasmodium parasites [8,9].

Basil genotypes have a wide variety of properties such as leaf sizes, color (green to dark purple), flower color (white, red, lavender and purple), growth characteristics (shape, height and flowering time) and aroma [10]. Therefore, the standard identifier list (UPOV 2003) has been used in the Guide for Conducting the Differences, Uniformity and Stability Tests (UPOV 2003; available) by the International Association for the Protection of New Plant Varieties (UPOV) to identify the varieties correctly. The UPOV system of plant variety protection based on an individual test guidelines represent an agreed and harmonized approach for the examination of new cultivars of a species of interest. Several studies deal with grouping of basil varieties based on only one nutritional characteristics according to chemotype [11,12], phenolic acid concentration [13,14] etc.

In recent years, the need for standard and quality materials in the sectoral demands of medicinal and aromatic plants has revealed the necessity of cultivating these plants and has also accelerated the variety development activities. Characterization of genetic resources has always remained one of the favorite methods of scientific community for the development of improved cultivars expressing higher yield with better quality and secondary metabolite. Knowledge of the genetic diversity in the germplasm populations is important for the efficient germplasm management and long-term breeding programs.

Industrialized countries can produce high-yield and quality products at low cost by using advanced breeding and agricultural techniques in the agriculture of medicinal and aromatic plants [15]. As in all agricultural products, medicinal and aromatic plants with economic importance are also preferred in certain standards. A good number of studies have been conducted to explain the morphological, phenological, and agronomic variability among local populations of sweet basil in diferent parts of the world [1,4,13,14,16]. In our country, Telci et al. [17] more than 80 local basil genotypes which obtained from different regions in Turkey determined the characterization of morphology, agronomic and technological. Up to now, earlier studies with medicine and aromatic plants have been conducted with local populations, it is necessary to investigate genetically and chemically important overseas genotypes in addition to targeted cultivation and breeding practices for desired morpho-chemotypes.

For this purpose, the present study was conducted in order to compare for the first genetic linkage map of 73 abroad, one local (Bolu) basil genotypes and one cultivar which is expected to facilitate the researches on genetics and breeding in basil cultivars regarding their agro-morphological traits as well as their essential oil yield, and to find the relationships likely to exist between morphological traits and essential oil constituents. Also, in this study, morphological and phenological features of genotypes which show superior characteristics in terms of yield will be determined by characterizing UPOV according to various evaluation criteria. Such information would be important to indicate the effect of geographic origin on agro-morphological and biochemical traits of basil seed cultivars. Promising basil cultivars can be used in various breeding programs and have the potential of enhancing its utilization.

## 2. Materials and Methods

### 2.1. Plant Material and Field Experiments

The experimental material comprised of seventyfour genotypes of basil (*Ocimum basilicum*) obtained from different geographical regions of the world (Supplementary Table 1). 73 basil genotypes, received from United States Department of Agriculture (USDA) with 4 commercial cultivars (Dino, midnigh, large sweet, moonlight) and local basil genotype were used in this study. These cultivars were developed through the single plant selection having resistance to various diseases and have been used as standard cultivars. All of them was grown in 2017 growing season, and except 14 genotypes (8 *Ocimum* × *africanum*, 3 *O. americanum and* 3 *O. basilicum*), adapted to Bolu ecological conditions (Table 1). In 2017, 59 genotypes, one local genotype and one cutivar were grown under field conditions in augmented design followed by selfing and single plant selection. Based on the adapted genotype, 50 genotypes were selected, and these genotypes were established in augmented design in 2018.

**Table 1.**
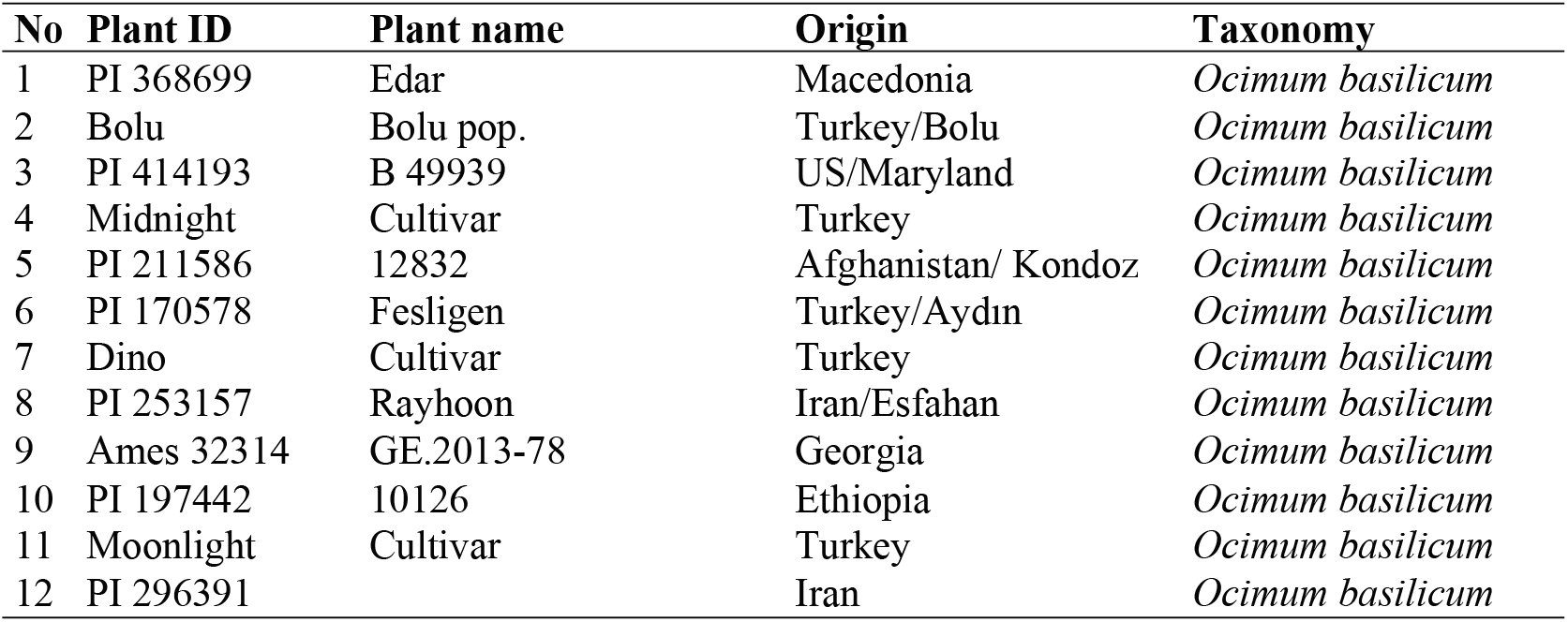
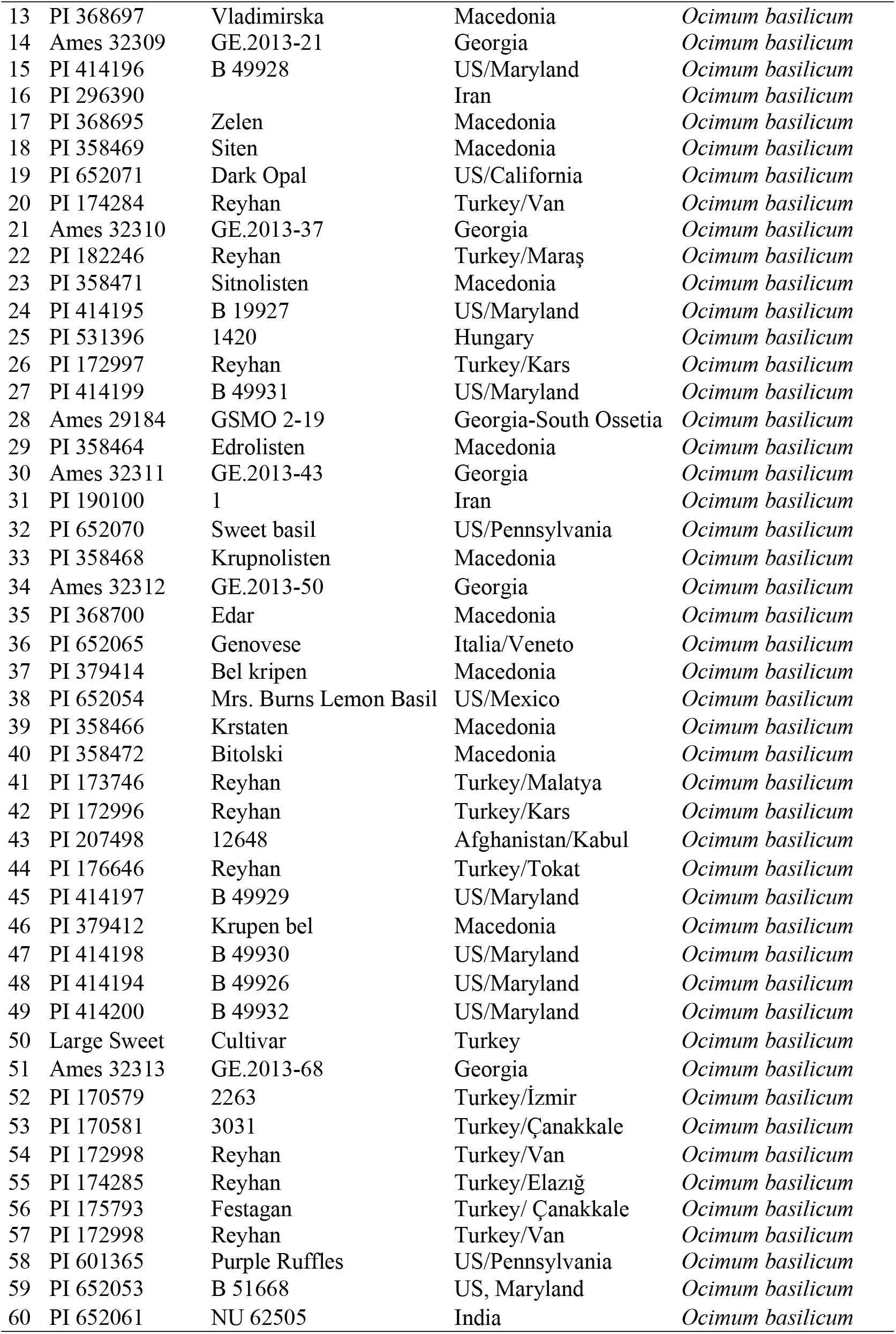
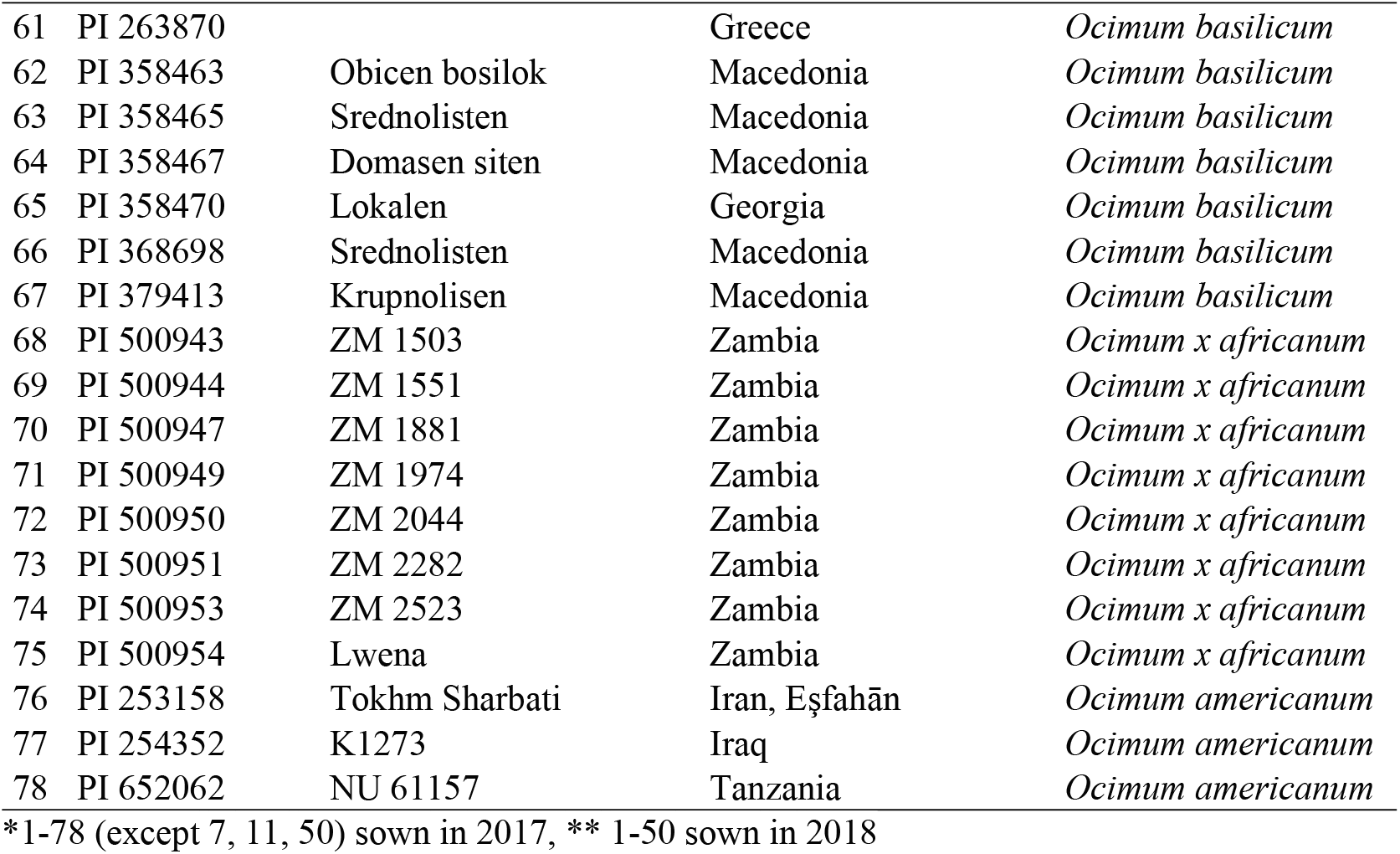
Plan ID, plant name and origin of basil genotypes.

The field experimental site was located at research and application area of Agriculture and Natural Sciences Faculty, is between 40°44’45’’N latitude, 3l°37’ 46”E longitudes with altitude of 752m. In first experimental year, since genotypes have very few seeds (up to 100), seeds were sown in a mixture of peat and perlite (9:1) on 19 april 2017. When the basil seedlings reached 10 cm in plant height, seedlings were transplanted into pilots on 15 May 2017 at a rate of 5 plants m^2^. In second experimental year, seeds were sown directly by hand on the field condition on 15 April 2018, each consisting of 4 m-long rows, row width and intra row spacing were 30 cm and 20 cm, respectively.

Average climatic data were recorded 16.08 °C temperature; 41.37 mm rainfall; 69.2% humidity during the vegetation period for 2017 and 17.10 °C temperature; 71.18 mm rainfall; 53.27% humidity in the growing season of 2018 [18]. Sweet basil was regularly irrigated to demonstrate good progress in its period vegetation since irrigation is a very important factor for cultivation of basil. As experimental factors, conventional fertilizer 60 kg ha^−1^ Diammonium phosphate (DAP) and 20 kg ha^−1^ Ammonium sulfate (AS) were applied with sowing all experimental years. Nitrogenous fertilizer as AS (in total 60 kg ha^−1^) was divided by two and applied to the plants in two splits in sown time and after first harvest of plant.

After each harvested, nitrogenous fertilizer as AS (in total 60 kg ha^−1^) was applied to the plants in two splits in July and August. Furthermore, when required, irrigation and weed control was made. Field data were collected by cutting randomly 10 plants from each plot, and the yield component of each plant was considered as the average for each plot in 2017 and 2018 years.

Plants were harvested by hand when the plants reached at the beginning of flowering, cutting off the overground part of the stem above its lignified fragments, and in two phases in total, 20-30 cm above the ground level during each growing season. The first harvest was at the beginning of flowering, and started from the first week of July until the third week of August in all experimental years. The second harvest was at the beginning of flowering, and started from the last week of July until the first week of October in all experimental years.

During harvest height of plants, as well as number of branchings in the first row of main stem were measured, as well as the weight of plant overground part. Then the herb was dried in thermal drying compartment in the temperature of 35°C and the air-dry herb weight was determined.

### 2.2. Morphological research and descriptor list

Morphological research was carried out during the year 2017 and 2018 in the field trial. Each genotypes was represented by 10 plants and total of 610 and 500 plants were analyzed for morphological traits in 2017 and 2018, respectively. Twenty-three traits were scored according to the Guidelines for the Conduct of Tests for Distinctness, Uniformity and Stability (UPOV 2003) in 2018 in which the qualitative traits were expressed in discontinuous states, while the expression of each quantitative trait was divided into a number of discrete states for the purpose of description (Table 2). All states are necessary to describe the full range of the traits, and every form of expression can be described by a single state. Thus, all the recorded data were qualitative in nature.

**Table 2.**
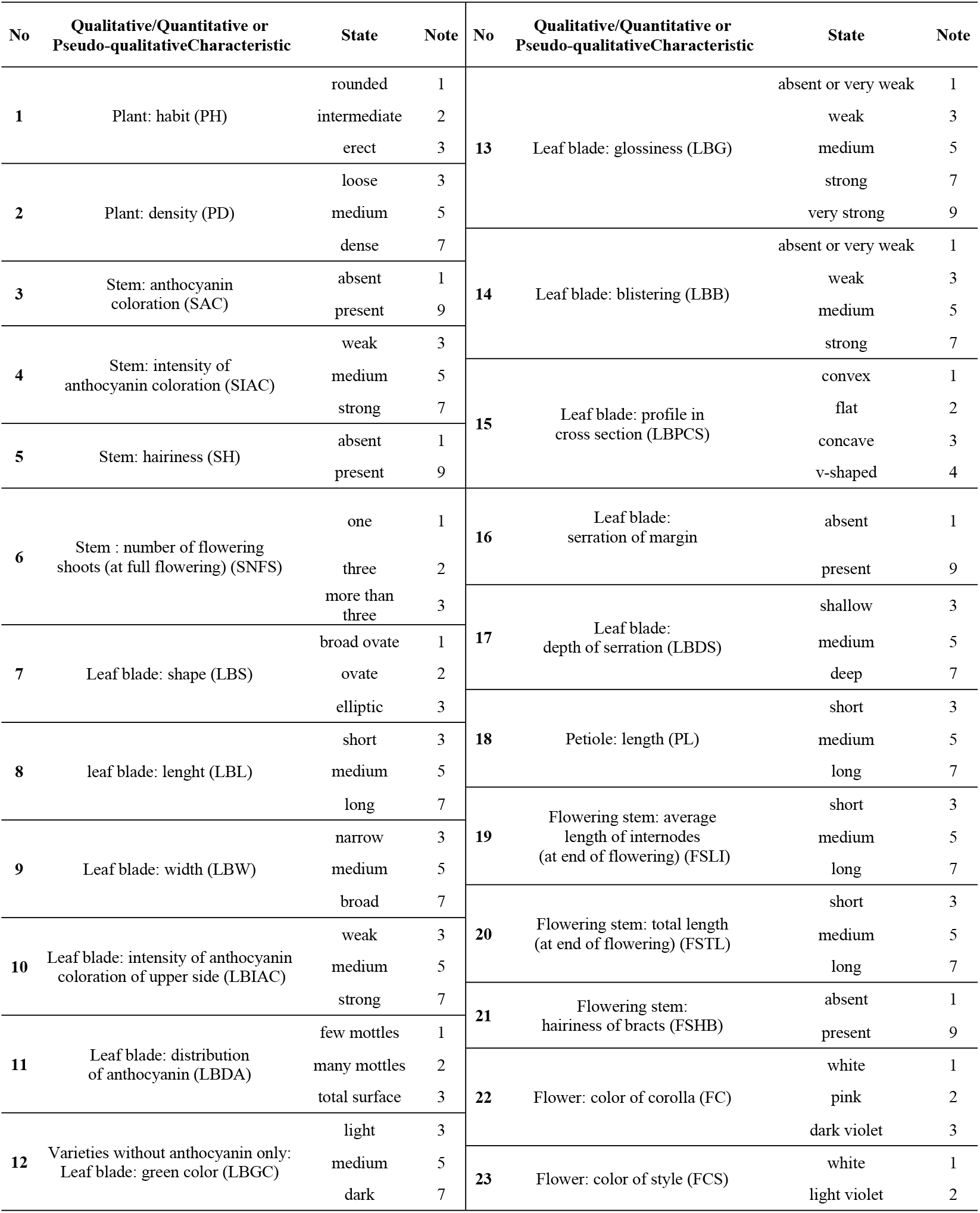
UPOV information of basil genotypes characteristic.

### 2.3. Essential oil extraction and analysis

A Clevenger device was used to obtain essential oil of basil by hydro-distillation. Fifty g of basil flowering aerial parts were taken from each treatment. Afterward, they were roughly crushed and placed in a 1-L glass balloon to which 500 mL of distilled water were added. Distillation was conducted for three h to obtain the essential oil content. Once collected in a sealed glass vials, the essential oil was dehydrated anhydrous sodium sulfate and stored at 4°C in darkness until analyses.

### 2.4. Data analysis

The differences among the basil genotypes were analysed through least significant differences (LSD) tests at 0.05 level of significance. Statistics included mean, range (minimum, maximum), coefficient of variation (CV) and standard deviations in 2017 and 2018 years. Cluster analysis was performed depending on the leaf shape and color of basil genotypes in 2017 and also, it was carried depending on the UPOV criteria among the basil genotypes using Ward’s method and squared Euclidian distance in 2018 experimental year [19,20]. Principal component analysis (PCA) and correlation analysis were conducted to determine the relationships among the morphological and yield properties of basil genotypes obtained in experimental years.

## 3. Results and Discussion

### 3.1 Observations

All phenotypic traits showed variation among the genotypes examined. Our results show that with a careful analysis and stringent selection of traits, morphological markers provide an inexpensive and reliable method for routine screening of a large number of genotypes, in order to monitor and manage germplasm collections.

The selection and choice of parents also depends upon the contribution of characters towards the genetic divergence (Table 3 and 4). Among the five characters studied, contribution percentage/fold was recorded maximum for branches/plant (27.69 %), followed by plant height (28.68%), days to 50% flowering (64.37%), oil content (about twenty fold), herb yield/plant (about eighteen fold) in 2017 experimental year, maximum for branches/plant (40.63 %), followed by plant height (32.7%), days to 50% flowering (73.63%), oil content (about twentyseven fold), in 2018 experimental year. Hence, these characters could be given due importance.

**Table 3.**
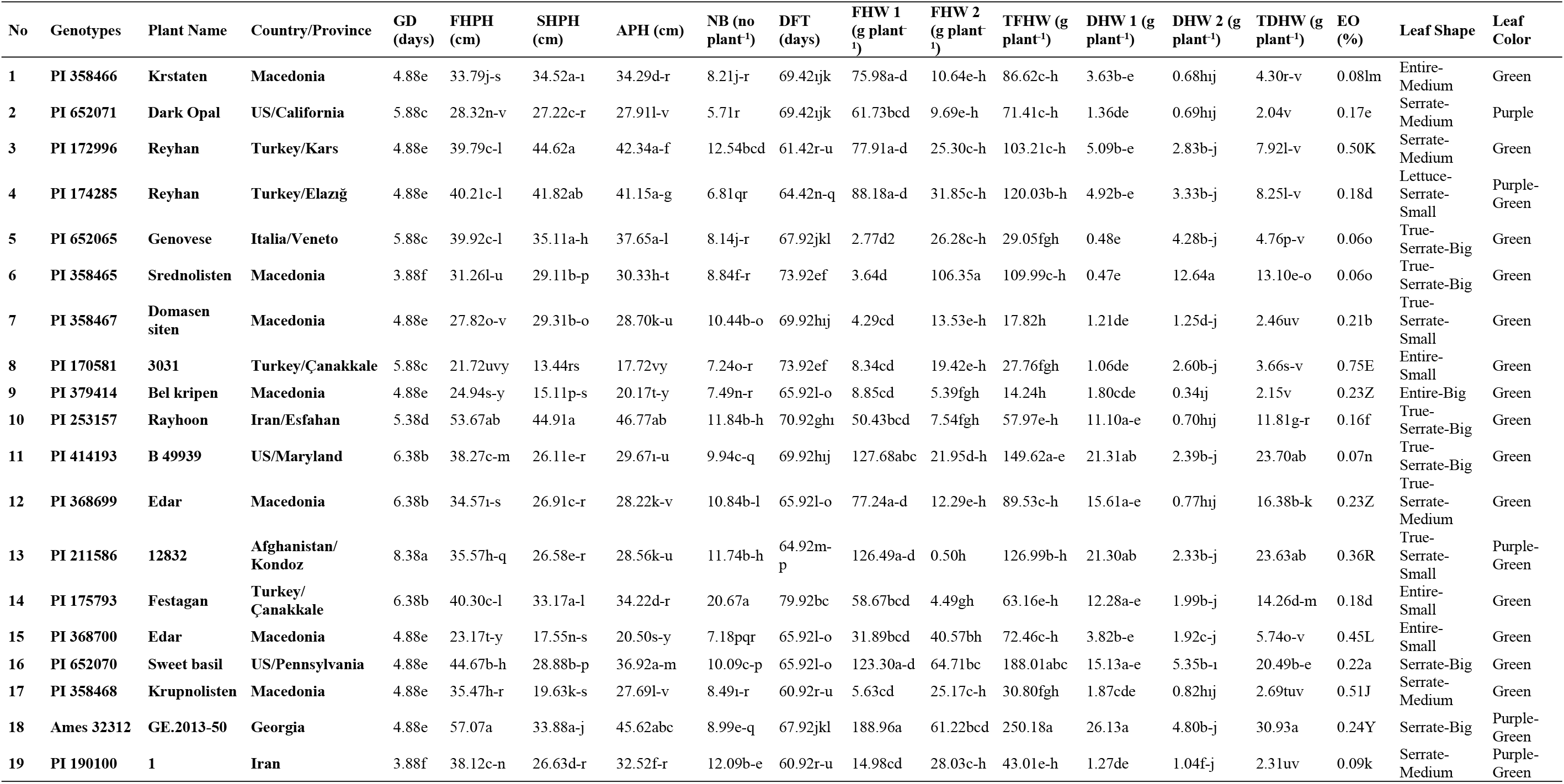
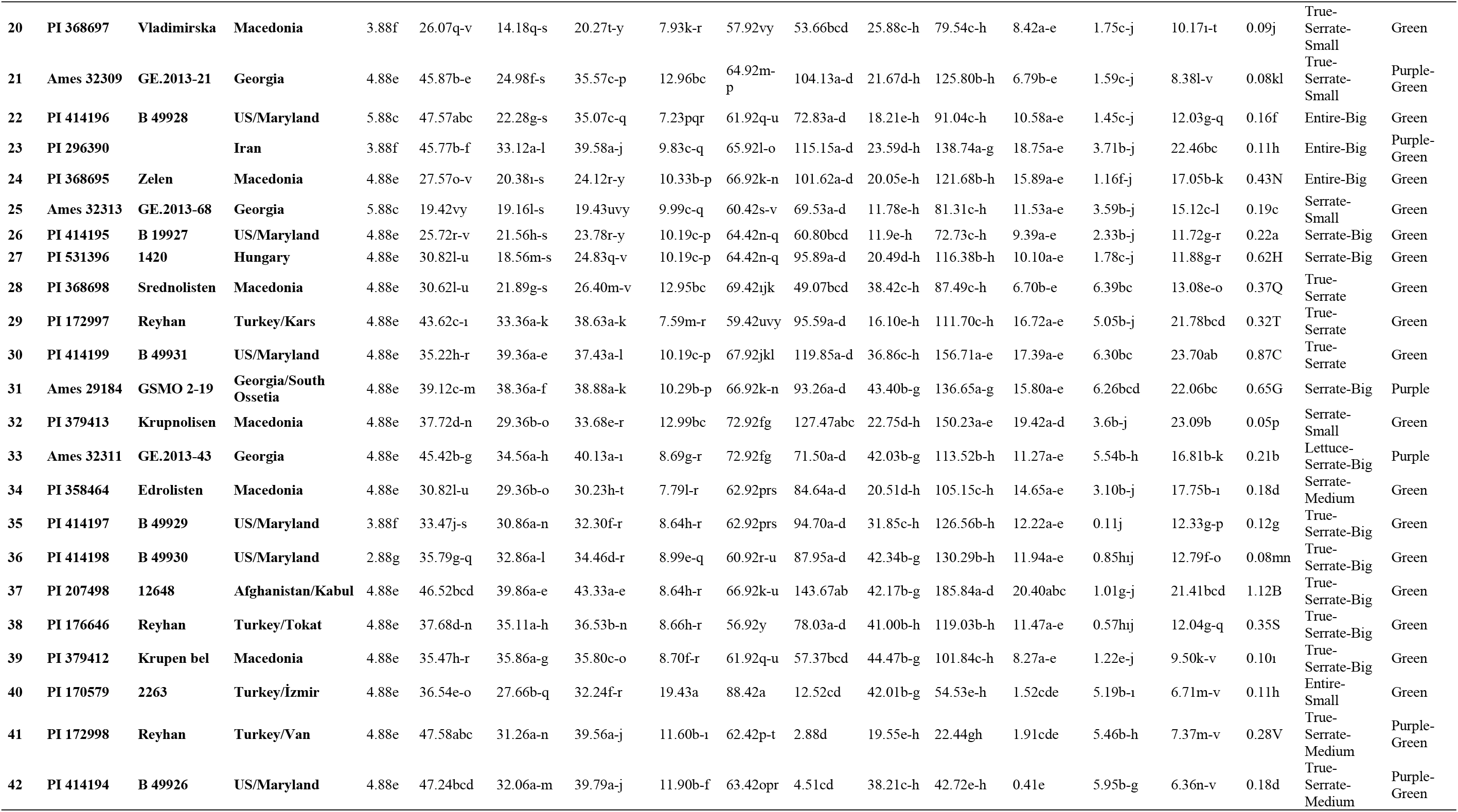
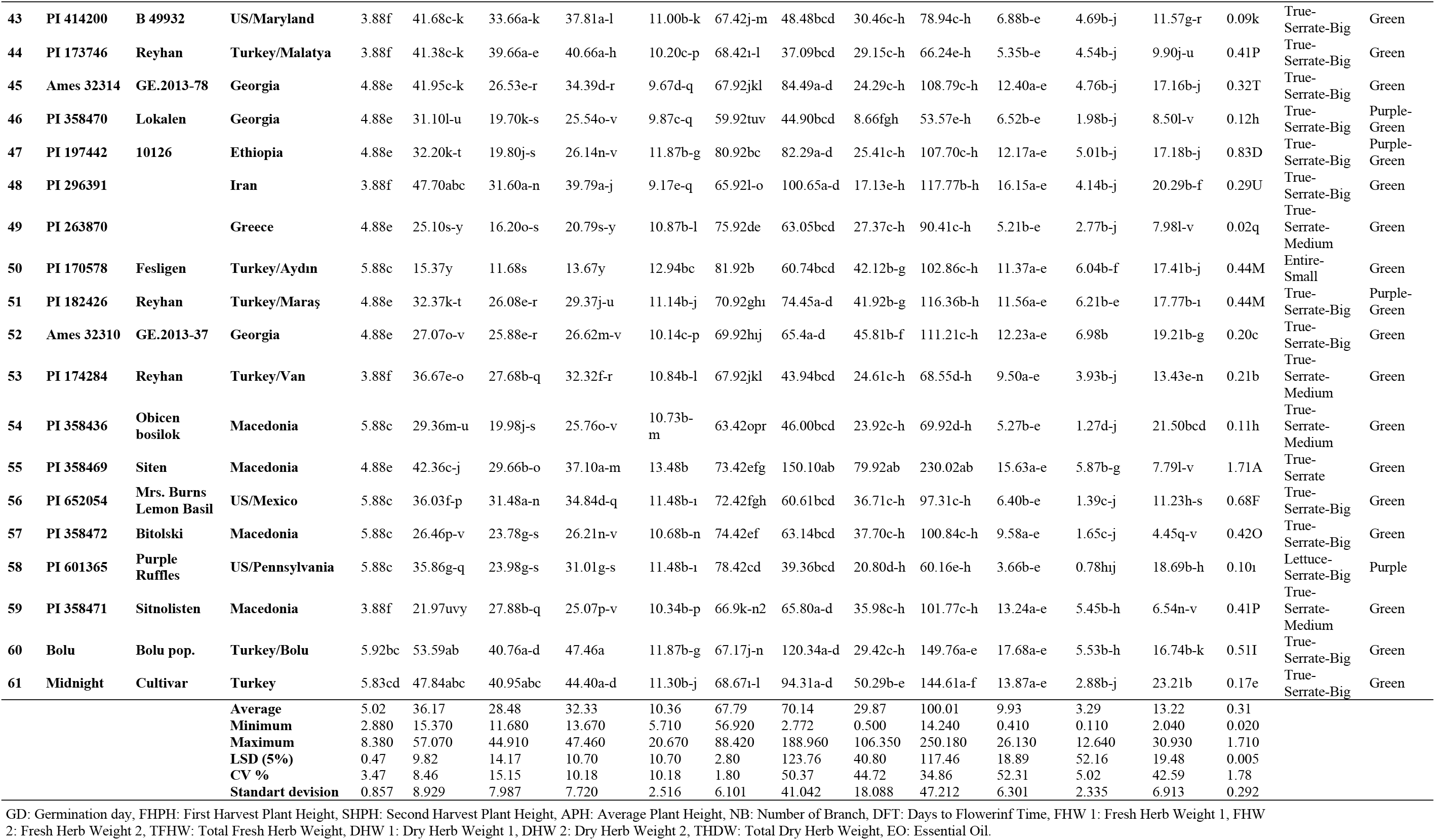
Morphological tarits of basil genotypes in 2017.

**Table 4.**
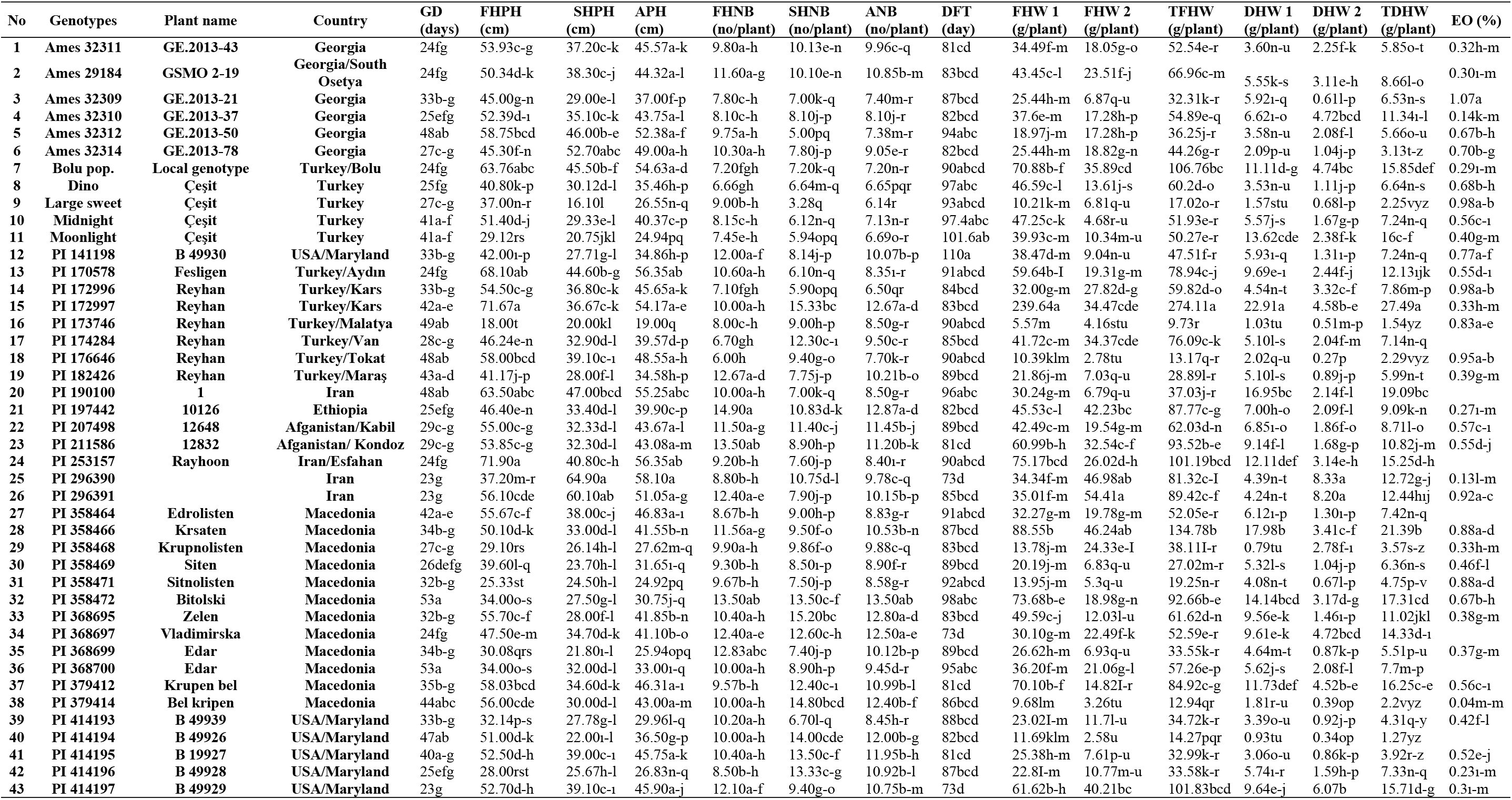
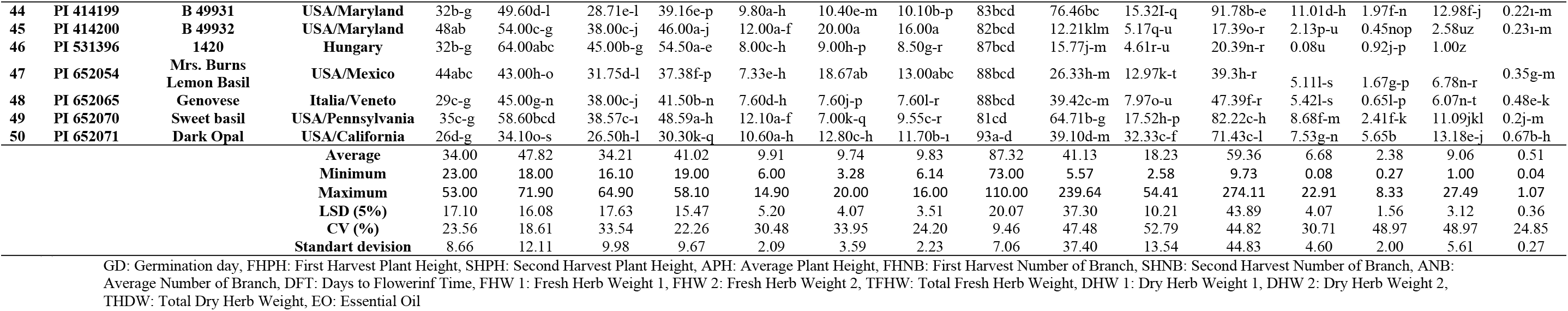
Morphological traits of basil genotypes in 2018.

Phenological observations of each genotypes were recorded, five a week. The number of days on which observations on seedling were made 2.88-8.38 in 2017, 23-53 in 2018. Observations on flowering day for first cutting were made 56.92-88.42 in 2017, 73-101.6 in 2018. During harvesting, the dates that basils were collected from each genotypes were recorded in experimantal years.

Our results were also in harmony with Srivastava et al. [21] noted that basil harvest takes about 85-90 days for maturity, when lower leaves start turning yellow and full blooming condition appears. Similarly, Bahl et al. [22] reported that three different basil genotypes (CIM-Saumya, CIM-Snigdha, CIM-Surabhi) matured in 80-100 days.

Leaf shape and leaf color were observed in 61 genotypes in 2017 year. Leaf shape of basil genotypes were divided to 12 different properties (4 entire-big, 1 entire-medium, 5 entire-small, 2 lettuce-serrate-big, 1 lettuce-serrate-small, 5 serrate-big, 5 serrate-medium, 2 serrate-small, 4 true-serrate, 21 true-serrate-big, 7 true-serrate-medium, 4 true-serrate-small). About 35% of basil genotypes had similar leaf shape being true-serrate big. Leaf colors of basil genotypes were found into three categories as green (46 genotypes), purple (4 genotypes) and purple-green (11 genotypes) in 2017 (Table 3).

In a study by Darrah [23], who classified the *O. basilicum* genotypes into seven categories: (1) tall slender types, (the sweet basil group); (2) large-leafed robust types (‘lettuc leaf’ also called ‘Italian’ basil); (3) dwarf types, which are short and small leafed (‘bush’ basil); (4) compact types, also described *O. basilicum* var. thyrsiflora (‘Thai’ basil); (5) purpurascens, the purple-colored basil types with traditional sweet basil flavor; (6) purple types (‘Dark Opal’: hybrid between *O. basilicum* and *O. forskolei* with a sweet basil plus clove-like aroma); and (7) citriodorum types (lemon-flavored basils).

Leaf shape and color of our basil genotypes were found similar with Darrah [25].

Egata et al. [24] recorded the morpho-agronomic variability to further select promising sweet basil accessions in Ethiopia. Singh et al. examined the genetic diversity and clustering pattern among 20 five basil accessions and found oil content was the highest contributing character toward the genetic diversity (56.09%). Srivastava et al. [21] studied 60 *O. basilicum* accessions and found some unique chemotypes which could be further exploited in future breeding programs. Carović-Stanko et al. [16] studied the resolving power of morphological traits for reliable identification of *O. basilicum* accessions and categorized six clusters of basil morphotypes.

### 3.2 Morphological parameters (plant height and number of branches)

The following morphological parameters were determined the plant height and number of branches. Plant height was determined two times before each sampling by measuring the height of ten randomly selected plants per plot from the soil to the top of the plant and getting an average value for each plot. These quantitative traits were investigated on the basis of the basil descriptors developed by the International Union for the Protection of New Varieties of Plants (UPOV, 2003) and were determined on ten randomly selected plants from the center rows of each cultivar. As can be seen in Table S3 and S4, significant differences were observed between the studied cultivars for growth parameters.

There were statistical significant differences among harvests for different genotypes with respect to the plant height in two experimental years. The presence of difference between the highest and the lowest values indicated that the genotypes included in the present study were quite diverse. In first year, while the highest plant height was obtained from Ames 32312 (57.07 cm) followed by PI 253157 (53.67 cm), and PI 172998 (47.58 cm) in first harvest, the highest plant height was obtained from PI 253157 (44.91 cm) followed by midnight (40.95 cm), and Bolu (40.76 cm) in second harvest. The highest avarage plant heights were recorded in Bolu (47.46 cm), PI 253157 (46.77 cm), Ames 32312 (45.62 cm), midnight (44.40 cm) genotypes. In second year, PI 253157 (71.90 cm) PI 170578 (68.10 cm) and PI 531396 (64.00 cm) genotypes have the highest plant height, whereas PI 173746 (18.00 cm), Moonlight (29.12 cm) have the lowest plant height in first harvest. In the second harvest, the highest plant height was recorded in PI 296390 (64.90 cm), followed by PI 296391 (60.10 cm) and Ames 32314 (52.70 cm) genotypes, while the lowest plant height was Large sweet (16.10 cm) and Moonlight (20.75 cm) cultivars. The highest avarage plant height was found at PI 296390 (58.10 cm) genotype, followed by PI 253157 (56.35 cm) and PI170578 (56.35 cm) genotypes. Especially, PI 253157 genotype have the highest plant height in experimental years.

The analysis of variance indicated the presence of considerable variability among the genotypes for the number of branches which was found to be highly significant (p=0.05) (Table 3). In addition, number of branches showed significant interaction of the different factors that were studied.

The number of branches was affected by genotypes, and by the interaction of year with genotype and harvest times. The genotype with the more branches was PI 175793 (20.67 number plant^−1^), followed by the PI 170579 (19.43 number plant^−1^), PI 358469 (13.48 number plant^−1^), Ames 32309 (12.96 number plant^−1^) and PI 368698 (12.95 number plant^−1^) genotypes. The genotype with the lowest number of branches was PI 652071(5.7 number/plant) followed by the PI 174285 (6.81 number plant^−1^) and PI 368700 (7.18 number plant^−1^) genotypes in first year. In the second year, the highest branch numbers were observed in PI 197442 (14.90 number plant^−1^), PI 211586 (13.50 number plant^−1^), PI 182426 (12.67 number plant^−1^) genotypes, the lowest branch numbers were observed in PI176646 (6.0 number plant^−1^), Dino (6.66 number plant^−1^), PI174284 (6.70 number plant^−1^) genotypes in first harvest. Moreover, the genotypes PI 414200 and PI 652054 showed the significantly and remarkably higher branch numbers and ranged between 20.00-18.67 number whereas, the lower one was observed in the registered genotype PI172996 (5.90 number) in second harvest. In addition, avarage the highest branch number was found at PI 414200 (16.00 number), followed by PI 358472 (13.50 number) and PI 652054 (13.00 number) genotypes.

The mean values of number of branches at the first year were higher than those of the second years, which was related to their height and habit, and all experimental years the highest branche numbers were obtained from first harvests. Based on our experimental results, there were significant differences between the two years of the experiments, and this can be because of the weather conditions as in 2018 the temperature was higher in vegetative growth period and the rainfall was much lower [18].

The examined basil cultivars had significant mean plant height (13.67-71.90 cm) in experimental years.

In earlier studies Egata et al. [24] reported that variability of combined analysis result on quantitative traits of Ethiopian sweet basil genotypes showed wide range in; number of primary branch/plant 6.08 to 8.98, number of internodes/main stem varies from 4.68 to 6.80, plant height 24.43 to 42.24 cm. Likewise, Alemu et al. [26] indicated that plant height was significantly affected by genotype, plant spacing, and they also reported that plant height changed from 52 to and 32 cm. Similarly, the plant height of different basil genotypes found at 20-60 cm [27], 22.9-57.0 cm [12], 40.0-76.9 cm [28], 60.89 cm [29], 65-88 cm [30], 69.7-89.5 cm [31] and 37.9-98.7 cm [32,33], 59.28-62.5 cm [34], 35.40-73.07 cm 35.40-73.07 cm [35], 24.43-42.24 cm [24], 17,16-45.33 cm [36], 74-80 cm [42] under different ecological conditions. Also, the morphological characters of 80 genotypes of *O. basilicum* from different geographical regions (India, Singapore, Tanzania, Thailand, and Slovak Republic) revealed a wide range of variation among themselves, viz. plant height (56-126 cm) [37–41].

The present results that we determined comply with the mentioned results with respect to plant height for two experimental years.

The examined basil cultivars had significant mean branch number (3.28-19.43 number/plant) in experimental years. These data corroborate with those reported by Egata et al. [24] (5.87-9.60 branches plant^−1^), 11.3-13.5 branches plant^−1^ [31], 10-12 branches plant^−1^ [30], 9.3 to 9.67 branches plant^−1^ [35], however, these data much lower than those of 8-41 branches plant^−1^ [42]. The number of branches was affected by the irrigation, cultivar, and the interaction of cultivar and year, growth stage and year, and the interaction of cultivar, year, and growth stage. Morphological characteristics, such as number of branches, are affected by irrigation, fertilization, and cultivar [43].

### 3.3 Fresh and dry weight

The analysis of variance revealed that there was an abundant scope for selection of promising lines from the present genetic stock for fresh and dry weight. The fresh and dry herb weights were changed by growth stages, year, and genotypes. The presence of huge difference between the highest and the lowest values indicated that the genotypes included in the present study were quite diverse in all experimental years.

In 2017, the genotypes were collected from Georgia (Ames 32312) and Macedonia (PI 358469) region and the character making it distinct was the high fresh yield plant^−1^ (250.18 g plant^−1^, 230.02 g plant^−1^, respectively), in contrast to other genotypes which showed fresh herb yield in the range of 14.24-156.71 g plant^−1^. Among the sixty genotypes, the highest dry weight was found Ames 32312 (30.93 g plant^−1^), followed by PI 414193 (23.70 g plant^−1^), PI 414199 (23.70 g plant^−1^), PI 211586 (23.63 g plant^−1^); however, the lowest dry weight was obtained from PI 652071 (2.04 g plant^−1^) and PI 379414 (2.15 g plant^−1^) genotypes.

In 2018, the highest fresh weight was obtained from PI 172997 (274.11 g plant^−1^), followed by PI358466 (134.78 g plant^−1^) and Bolu (106.76 g plant^−1^) genotypes, in contrast to other genotypes which showed herb yield in the range of 12.94-101,83 g plant^−1^ in 2018 experimantal year. The lowest was found at PI 379414(12.94 g plant^−1^) and PI 176646 (13.17 g plant^−1^) genotypes The genotype that showed the highest dry weight was PI 172997 (27.49 g plant^−1^), followed by PI358466 (21.39 g plant^−1^) and PI379412 (16.25 g plant^−1^) in second years; however, the lowest dry weight was found PI 531396 (1.0 g plant^−1^) and PI 414194 (1.27 g plant^−1^) genotypes. Most of the genotypes had yields greater than 50 g plant^−1^ fresh herbage, which are considered very high. Our results suggest that most basil genotypes can provide high yields under Bolu ecological condition. Especially, PI 172997, Ames 32312, PI 358469 genotypes were foremost in experimental years.

The examined basil cultivars had significant fresh herb yields in first experimental year (17.18-250.18 g/plant) and in second experimental year (12.94-274.11 g/plant) (Table 3 and 4). The examined basil cultivars had significant dry herb yields in first experimental year (2.04-30.93 g/plant) and in second experimental year (1.0-27.49 g/plant) (Table 3 and 4).

The results of the present study are in a good agreement with those of the study by Egata et al. [24], who reported that fresh leaf weight/plant varied from 44.58 g to 231.53 g, and also noted that maximum dry weight/plant ranged from 10.06 to 31.57 g in Ethiopian sweet basil genotypes. Likewise, previous studies have reported a range of values for fresh weight of different basil cultivars from 240.2 to1105.9 g m^−2^ and dry weight was in the range of 47.9-202.8 g m^−2^ [44–46].

Similarly, dry herb weight plant^−1^ varied from 7.26-10.78 g under stress condition [47], and ranged from 5.5-7.1 g grown with different temperature integration doses [48]. Our results agree with those obtained by previous literature.

By contrast, Kalamartzis et al. [49] determined the effect of water stress on five cultivars of basil (Mrs Burns, Cinnamon, Sweet, Red Rubin, Thai) were found to be within the ranges of 378.5-4357.5 g m^−2^ for fresh herb, 65.8-922.5 g m^−2^ for dry herb. In addition, Karaca et al. [36] determined that fresh herb yield and dry herb yield of nine basil genotypes were found to be within the ranges of 195.00-383.99 g plant^−1^, 22.21-46.85 g plant^−1^, respectively. Köse et al. [50] also, indicated that effect of plant density on fresh herbage yield of sweet basil was in the range of 192.00-464.70 g plant^−1^ and dry weight was in range 52.90-9.98 g plant^−1^. Furthermore, Arslan et al. [51] noted the fresh and dry yield of *O. basilicum* in the range of 192.00-464.70, also 24.3-55.2 g under the Eastern Mediterranean condition, respectively.

The observed differences in the fresh and dry plant herb yield across countries may be a result of different environmental and genetic factors, different chemotypes, harvest time, weather conditions and the cultural practices.

### 3.4 Essential oil content

Essential oil content was affected by growth stages, year, cultivar, and also by their interactions (Table 3, 4). It was found that the significant differences were seen among the used fennel genotypes in terms of essential oil content at p<0.05 (Table 3). The highest essential oil content was found at PI 358469 (1.71 %), followed by PI 207498 (1.12%), PI 414199 (0.87%), PI 197442 (0.83 %) in first experimental year. The lowest essential oil content was found at PI 379413 (0.05 %) (Table 3 and 4).

The highest essential oil content was found at Ames 32309 (1.07 %), followed by PI 172996 (0.98%) and large sweet (0.98%) in second years. The lowest essential oil content was found at PI 379414 (0.04 %) and PI 296390 (0.13%) genotypes.

Great variations in the essential oil content of *O. basilicum* across geographic regions might be attributed to variable agroclimatic conditions and different agronomic techniques for cultivating [52,53]. The essential oil yield (0.04-1.71%) in the present study was comparable to that in a study by Egata et al. [24], who found the range of 0.10-1.02 % leaf essential oil content dry weight base for Ethiopian sweet basil genotypes. Similarly, Karaca et al. [36] determined that essential oil content of in nine basil genotypes were found to be within the ranges of 0.25-1.06%. Also Beatovićr et al. [54] reported that the essential oil yields of twelve *Ocimum basilicum* L. cultivars grown in Serbia ranged from 0.65 to 1.90 %.

Likewise, Telci et al. [12] reported values for essential oil content between 18 genotypes ranging from 0.4% to 1.5%. Zheljazkov et al. [4] recorded that essential oil content of the tested 38 genotypes a grown in Mississippi varied from 0.07% to 1.92% in dry herbage in the field experiment. It was reported that essential oil yield of the air-dried overground parts of *Ocimum basilicum* from Turkey as obtained by hydrodistillation was 1.25% [55]. Juliani et al. [56] reported basil oil content from 0.6 to 1.7% dry herbage. The yield of essential oil from different plant parts varies between 0.15-1.59%, and it depends also on the seasonal factor and locality [57–62].

By contrast, Simon et al. [5] reported the essential oil content from 0.04% to 0.70% (v fresh weight^−1^) in a study of a large number of basil genotypes. In addition, the content and composition of the essential oil have been evaluated in 14 *O. basilicum* genotypes, and reported that varied from 0.59 to 2.30% (genotype no. 6) [63]. Furthermore, in four Ocimum species grown in Tanzania (Runyoro et al., 2010), the essential oil yields ranged from 0.5% to 4%.

According to our study, the USDA genotypes were higher in its content of essential oil in compare with the local genotypes. Based on essential oil content (Table 4), PI 358469 originating from Macedonia and Ames 32309 originating from Georgia are the best genotypes, which are the different from the other genotypes.

In general, the essential oil content of basil genotypes in this study was within the usual content reported in other studies [4,24,64]; however, Runyoro et al. [65] and Akçali Giachino et al. [63] results that were higher than our results. Differences in basil essential oil content between this study and another report from research could be due to differential environmental conditions, their genetics and growth conditions [66,67].

### 3.5 Correlation analysis and principal component analysis

Correlation analysis assists in selecting the effective properties in order to an indirect selection of superior genotypes. In addition to this, principal component analysis is an appropriate multivariate technique to identify and assessment of independent principle components depending on influential plant characteristics. These two analysis methods contribute to the plant breeding program to help breeders [68,69].

#### 3.5.1 Correlation analysis of basil genotypes

The assessment of correlation analysis values among the all examined properties for 2017 and 2018 years were shown in table 5 and table 6. This analysis was conducted to determine the relationship among the examined properties in 2017 and 2018 years. There was found 27 positives and one negative correlations totally which out of the 17 highly significant and positive correlations were observed between r=0.328-0.926, 8 positive correlations were seen between r= 0.273-0.320 and only one negative correlation was found between GD and FHW 2 with r=-0.325 in the 2017 year (Table 5). The highest positive correlation was noted between the PHPH and APH with r=0.926. Other high positive correlations were found between the FHPH and SHPH, FHW 1, TFHW and TDHW. Similarly, SHPH had positive correlation with APH. APH had recorded positive correlated with FHW 1 and TFHW. NB had only one correlation with DFT (r=0.547). FHW 1 correlated with TFHW, DHW and TDHW. FHW2 correlated with TFHW, DHW 2 and TFHW correlated with DHW 1, TDHW and EO. DHW1 had only one correlation with TDHW (r=0.822). Positive correlations also were obtained between the FHPH and DHW 1, and SHPH was found positive correlation with FHW1 and TFHW. DFT was found correlation with DHW 2. FHW 1 and FHW 2 had correlation with EO and DHW2 correlated with TDHW (r=0.274). EO of basil genotypes was correlated only fresh weight among the examined properties and most of correlations were found in yield properties fresh and dry weight in 2017 year.

**Table 5.** Correlation analysis among the examined properties of basil genotypes in 2017.

|  | GD<br>(days) | FHPH<br>(cm) | SHPH<br>(cm) | APH<br>(cm) | NB (no<br>plant<sup>-1</sup>) | DFT<br>(days) | FHW 1 (g<br>plant<sup>-1</sup>) | FHW 2 (g<br>plant<sup>-1</sup>) | TFHW (g<br>plant<sup>-1</sup>) | DHW 1 (g<br>plant<sup>-1</sup>) | DHW 2 (g<br>plant<sup>-1</sup>) | TDHW (g<br>plant<sup>-1</sup>) | EO (%) |
| --- | --- | --- | --- | --- | --- | --- | --- | --- | --- | --- | --- | --- | --- |
| GD (days) | 1 | -0.045 | -0.116 | -0.13 | 0.178 | 0.224 | 0.101 | -0.325* | -0.037 | 0.143 | -0.193 | 0.17 | 0.075 |
| FHPH (cm) | -0.045 | 1 | 0.703** | 0.926** | 0.126 | -0.128 | 0.368** | 0.123 | 0.367** | 0.32* | 0.097 | 0.328** | 0.008 |
| SHPH (cm) | -0.116 | 0.703** | 1 | 0.91** | 0.068 | -0.116 | 0.285* | 0.176 | 0.315* | 0.234 | 0.128 | 0.209 | 0.049 |
| APH (cm) | -0.13 | 0.926** | 0.91** | 1 | 0.083 | -0.138 | 0.348** | 0.2 | 0.379** | 0.274* | 0.139 | 0.277* | 0.051 |
| NB (no plant<sup>-1</sup>) | 0.178 | 0.126 | 0.068 | 0.083 | 1 | 0.547** | -0.019 | 0.01 | -0.013 | 0.045 | 0.167 | 0.076 | 0.063 |
| DFT (days) | 0.224 | -0.128 | -0.116 | -0.138 | 0.547** | 1 | -0.076 | 0.188 | 0.006 | -0.06 | 0.293* | 0.044 | 0.148 |
| FHW 1 (g plant<sup>-1</sup>) | 0.101 | 0.368** | 0.285* | 0.348** | -0.019 | -0.076 | 1 | 0.146 | 0.925** | 0.888** | 0.012 | 0.696** | 0.312* |
| FHW 2 (g plant<sup>-1</sup>) | -0.325* | 0.123 | 0.176 | 0.2 | 0.01 | 0.188 | 0.146 | 1 | 0.51** | 0.069 | 0.617** | 0.155 | 0.273* |
| TFHW (g plant<sup>-1</sup>) | -0.037 | 0.367** | 0.315* | 0.379** | -0.013 | 0.006 | 0.925** | 0.51** | 1 | 0.799** | 0.246 | 0.665** | 0.376** |
| DHW 1 (g plant<sup>-1</sup>) | 0.143 | 0.32* | 0.234 | 0.274* | 0.045 | -0.06 | 0.888** | 0.069 | 0.799** | 1 | 0.089 | 0.822** | 0.244 |
| DHW 2 (g plant<sup>-1</sup>) | -0.193 | 0.097 | 0.128 | 0.139 | 0.167 | 0.293* | 0.012 | 0.617** | 0.246 | 0.089 | 1 | 0.274* | 0.155 |
| TDHW (g plant<sup>-1</sup>) | 0.17 | 0.328** | 0.209 | 0.277* | 0.076 | 0.044 | 0.696** | 0.155 | 0.665** | 0.822** | 0.274* | 1 | 0.043 |
| EO (%) | 0.075 | 0.008 | 0.049 | 0.051 | 0.063 | 0.148 | 0.312* | 0.273* | 0.376** | 0.244 | 0.155 | 0.043 | 1 |

\*5%, \*\*1%; GD: Germination day, FHPH: First Harvest Plant Height, SHPH: Second Harvest Plant Height, APH: Average Plant Height, NB: Number of Branch, DFT: Days to Flowerinf Time, FHW 1: Fresh Herb Weight 1, FHW 2: Fresh Herb Weight 2, TFHW: Total Fresh Herb Weight, DHW 1: Dry Herb Weight 1, DHW 2: Dry Herb Weight 2, THDW: Total Dry Herb Weight, EO: Essential Oil

**Table 6.**
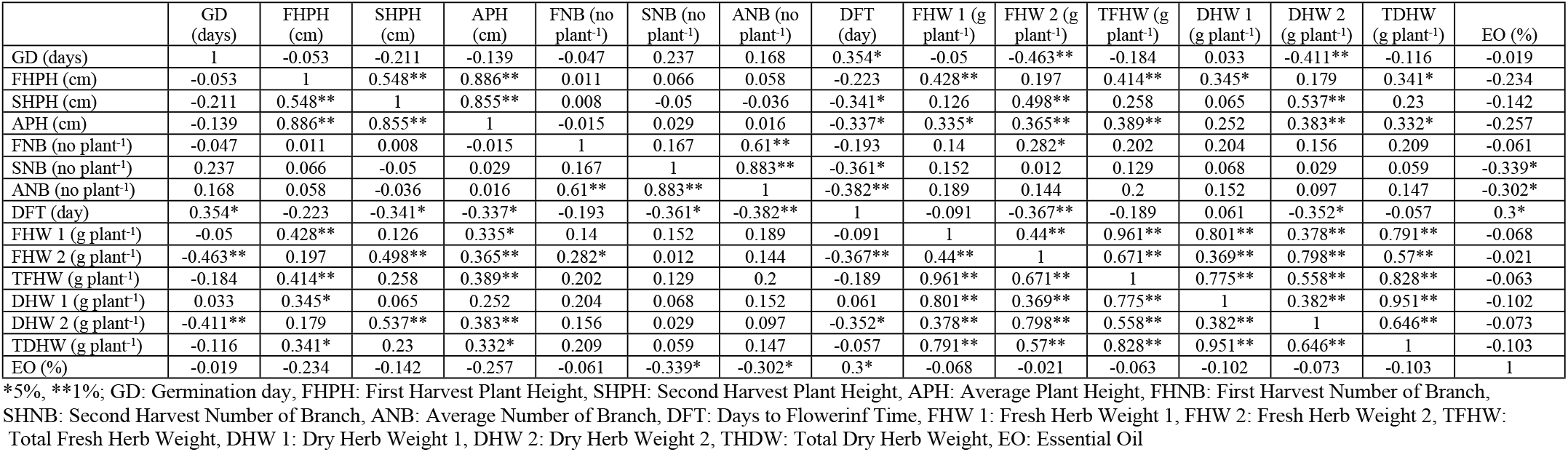
Correlation analysis among the examined properties of basil genotypes in 2018.

|  | GD<br>(days) | FHPH<br>(cm) | SHPH<br>(cm) | APH<br>(cm) | FNB (no<br>plant<sup>-1</sup>) | SNB (no<br>plant<sup>-1</sup>) | ANB (no<br>plant<sup>-1</sup>) | DFT<br>(day) | FHW 1 (g<br>plant<sup>-1</sup>) | FHW 2 (g<br>plant<sup>-1</sup>) | TFHW (g<br>plant<sup>-1</sup>) | DHW 1<br>(g plant<sup>-1</sup>) | DHW 2<br>(g plant<sup>-1</sup>) | TDHW<br>(g plant<sup>-1</sup>) | EO (%) |
| --- | --- | --- | --- | --- | --- | --- | --- | --- | --- | --- | --- | --- | --- | --- | --- |
| GD (days) | 1 | -0.053 | -0.211 | -0.139 | -0.047 | 0.237 | 0.168 | 0.354* | -0.05 | -0.463** | -0.184 | 0.033 | -0.411** | -0.116 | -0.019 |
| FHPH (cm) | -0.053 | 1 | 0.548** | 0.886** | 0.011 | 0.066 | 0.058 | -0.223 | 0.428** | 0.197 | 0.414** | 0.345* | 0.179 | 0.341* | -0.234 |
| SHPH (cm) | -0.211 | 0.548** | 1 | 0.855** | 0.008 | -0.05 | -0.036 | -0.341* | 0.126 | 0.498** | 0.258 | 0.065 | 0.537** | 0.23 | -0.142 |
| APH (cm) | -0.139 | 0.886** | 0.855** | 1 | -0.015 | 0.029 | 0.016 | -0.337* | 0.335* | 0.365** | 0.389** | 0.252 | 0.383** | 0.332* | -0.257 |
| FNB (no plant<sup>-1</sup>) | -0.047 | 0.011 | 0.008 | -0.015 | 1 | 0.167 | 0.61** | -0.193 | 0.14 | 0.282* | 0.202 | 0.204 | 0.156 | 0.209 | -0.061 |
| SNB (no plant<sup>-1</sup>) | 0.237 | 0.066 | -0.05 | 0.029 | 0.167 | 1 | 0.883** | -0.361* | 0.152 | 0.012 | 0.129 | 0.068 | 0.029 | 0.059 | -0.339* |
| ANB (no plant<sup>-1</sup>) | 0.168 | 0.058 | -0.036 | 0.016 | 0.61** | 0.883** | 1 | -0.382** | 0.189 | 0.144 | 0.2 | 0.152 | 0.097 | 0.147 | -0.302* |
| DFT (day) | 0.354* | -0.223 | -0.341* | -0.337* | -0.193 | -0.361* | -0.382** | 1 | -0.091 | -0.367** | -0.189 | 0.061 | -0.352* | -0.057 | 0.3* |
| FHW 1 (g plant<sup>-1</sup>) | -0.05 | 0.428** | 0.126 | 0.335* | 0.14 | 0.152 | 0.189 | -0.091 | 1 | 0.44** | 0.961** | 0.801** | 0.378** | 0.791** | -0.068 |
| FHW 2 (g plant<sup>-1</sup>) | -0.463** | 0.197 | 0.498** | 0.365** | 0.282* | 0.012 | 0.144 | -0.367** | 0.44** | 1 | 0.671** | 0.369** | 0.798** | 0.57** | -0.021 |
| TFHW (g plant<sup>-1</sup>) | -0.184 | 0.414** | 0.258 | 0.389** | 0.202 | 0.129 | 0.2 | -0.189 | 0.961** | 0.671** | 1 | 0.775** | 0.558** | 0.828** | -0.063 |
| DHW 1 (g plant<sup>-1</sup>) | 0.033 | 0.345* | 0.065 | 0.252 | 0.204 | 0.068 | 0.152 | 0.061 | 0.801** | 0.369** | 0.775** | 1 | 0.382** | 0.951** | -0.102 |
| DHW 2 (g plant<sup>-1</sup>) | -0.411** | 0.179 | 0.537** | 0.383** | 0.156 | 0.029 | 0.097 | -0.352* | 0.378** | 0.798** | 0.558** | 0.382** | 1 | 0.646** | -0.073 |
| TDHW (g plant<sup>-1</sup>) | -0.116 | 0.341* | 0.23 | 0.332* | 0.209 | 0.059 | 0.147 | -0.057 | 0.791** | 0.57** | 0.828** | 0.951** | 0.646** | 1 | -0.103 |
| EO (%) | -0.019 | -0.234 | -0.142 | -0.257 | -0.061 | -0.339* | -0.302* | 0.3* | -0.068 | -0.021 | -0.063 | -0.102 | -0.073 | -0.103 | 1 |

\*5%, \*\*1%; GD: Germination day, FHPH: First Harvest Plant Height, SHPH: Second Harvest Plant Height, APH: Average Plant Height, FHNB: First Harvest Number of Branch, SHNB: Second Harvest Number of Branch, ANB: Average Number of Branch, DFT: Days to Flowerinf Time, FHW 1: Fresh Herb Weight 1, FHW 2: Fresh Herb Weight 2, TFHW: Total Fresh Herb Weight, DHW 1: Dry Herb Weight 1, DHW 2: Dry Herb Weight 2, THDW: Total Dry Herb Weight, EO: Essential Oil

Forty-three correlations were found among the 15 examined properties in fifty basil genotypes as positive and negative in 2018 year. 26 high significant positive, 4 high significant negative, 7 positive and 6 negative correlations were obtained in correlation analysis of 2018 year. According to GD headings, data had a highly significant negative correlation with FHW 2 (r=-0.463) and DHW2 (r=-0.411) and positive correlation with DFT (r=0.354). Concerning to FHPH, data showed highly significant positive correlation with SHPH, APH, FHW 1 and TFHW with r=0.548, 0.886, 0.428, 0.414, respectively. It also gave positive correlation with DHW (r=0.345) and TDHW (r=0.341). As for SHPH, it showed highly significant positive correlation coefficient with APH (r=0.855), FHW 2 (r=0.498) and DHW 2 (r=0.537) while recorded negative correlation with DFT (r=0.341). Regarding APH, it showed a correlation with DFT (r=-0.337) as negatively and FHW 1 (r=0.335) and TDHW (r=0.332) as positively. Moreover, it gave highly significant positive correlation with FHW 2 (r=0.365), TFHW (r=0.389) and DHW 2 (r=0.383). FNB had one highly significant positive correlation with ANB (r=0.61) and one positive correlation with FHW2 with r=0.282. Although SNB showed a negative correlation with DFT and EO with r=-0.361 and r=-0.339, respectively, it had highly significant positive correlation with ANB (r=0.883). ANB had highly significant negative correlation with DFT (r=-0.382) and negative correlation with EO (r=-0.302). DFT had highly significant negative correlation with FHW 2 (r=-0.367) and negative correlation with DHW2 (r=-0.352). It also showed positive correlation coefficient with EO (r=0.3). Highly significant positive correlations were obtained for FHW 1 with FHW 2, TFHW, DHW 1, DHW 2 and TDHW with r=0.44, 0.961, 0.801, 0.378 and 0.791, respectively. FHW 2 had highly significant positive correlation with other yield values as TFHW (r=0.671), DHW 1 (r=0.369), DHW 2 (r=0.798) and TDHW (r=0.57). TFHW showed highly significant and positive correlation with DHW 1, DHW 2 and TDHW with r=0.775, 0.558 and 0.828, respectively. DHW 1 had highly significant positive correlation with DHW2 (r=0.382) and TDHW (r=0.951). At the same time, DHW 2 was also recorded as highly significant positive correlation with TDHW (r=0.646).

#### 3.5.2 Principal component analysis (PCA) of basil genotypes

PCA analysis carried out to separate the number of highly diverse genotypes of basil to support the breeders in crossing and selection breeding successful programs. Thus, prominent genotypes can be used as good parents in development programs of basil [70].

All basil morphological properties were evaluated together with in figure 1a and 1b for 2017 and 2018 years, respectively. PCA showed PC1 accounted for 33.97% of variation with GD, NB, DFT, FHW1, FHW2, DHW1, DHW2, TFHW and EO in 2017 year. FHW1, FHW2 and TFHW were the major factors for PC1. These properties were found closely on the same side of loading factor. PC2 showed 16.28% as an accounted and it had FHPH, SHPH and APH properties. PHPH was the major factors for PC2. PCA had 52.58% as totally in 2018. PC1 accounted for 35.99% of variation with FHW1 and DHW1 being the major factors. PC2 accounted for 16.59% variation with GD being the major contributor. FNB, SNB and ANB located on the same side of plot loading and DFT and EO located opposite side of these properties.

**Figure 1.**
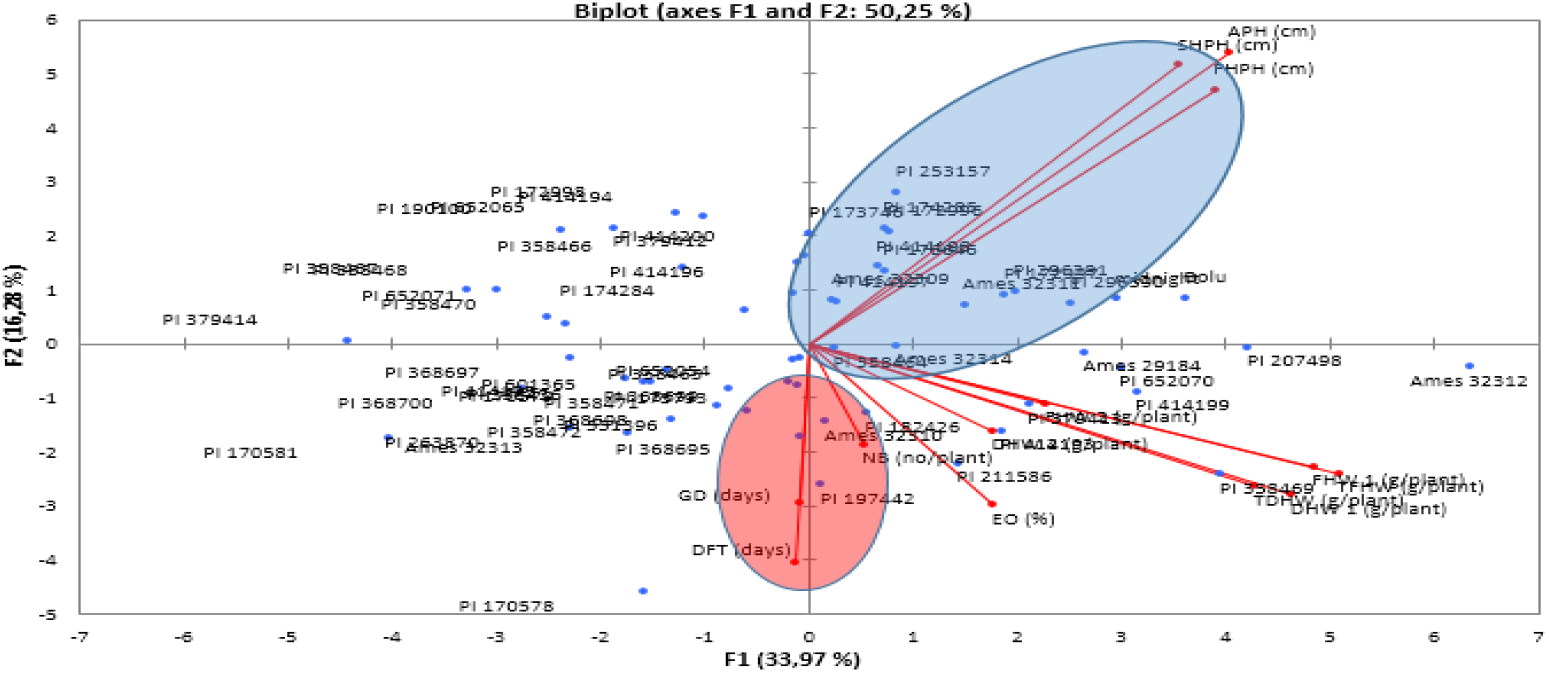
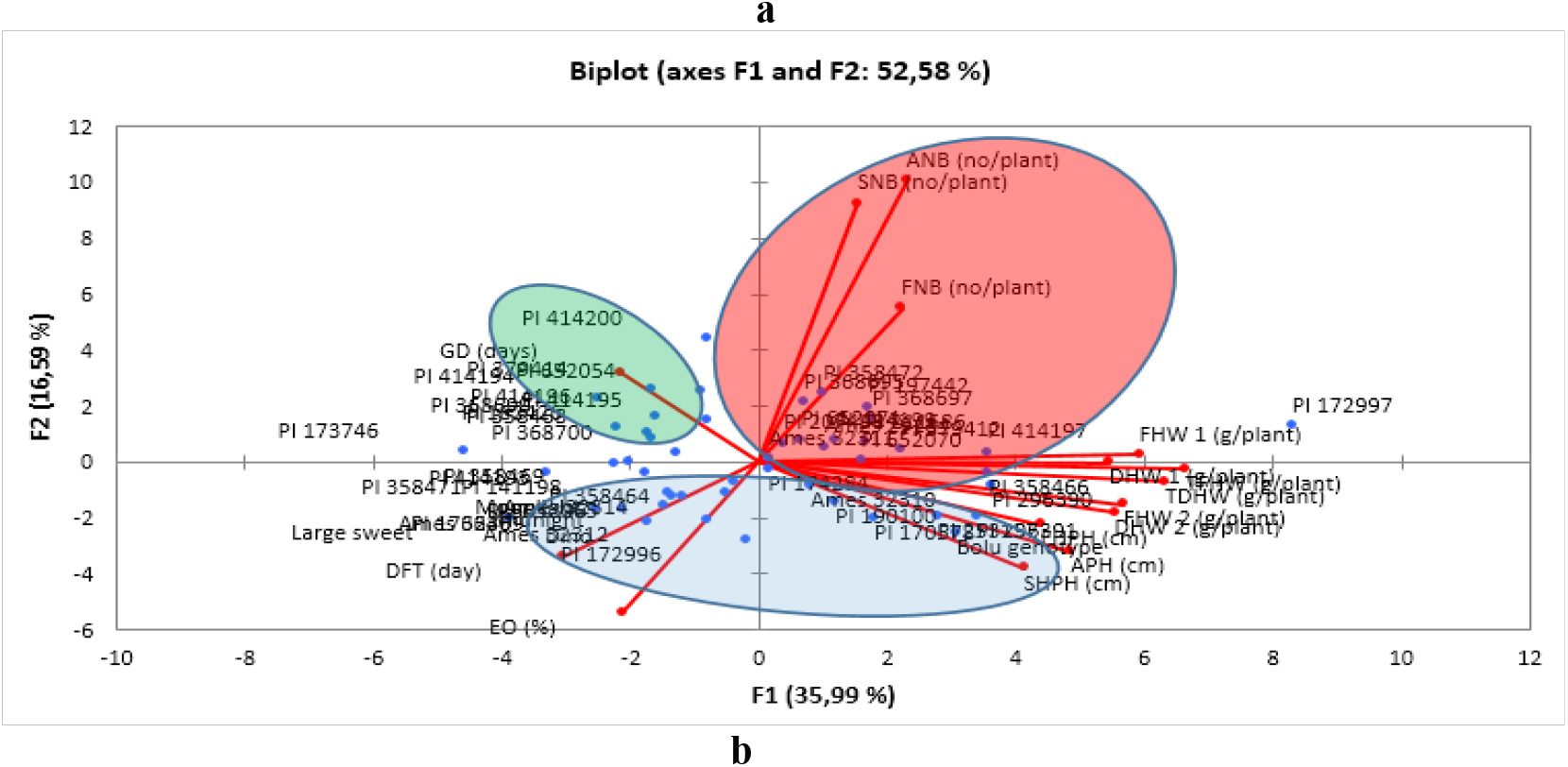
PCA analysis of basil genotypes in 2017 (a) and 2018 (b).

#### 3.5.3 Genetic diversity of basil genotypes

It was reported that plant properties depending on UPOV descriptor are highly heritable and reliable for the first classification of genotypes into different morphotypes. Morphological markers ensure a simple and cheap method for routine screening of many of genotypes to identify morphotypes and manage the germplasm collection of plants [16]. For this reason, a hierarchical cluster analysis (HCA) was performed to get the best classification depending on leaf color and leaf shape for 2017 and UPOV criteria by using the 2018 data. We selected the agglomeration method by using the Ward method for 2018. Constellation plot provided the best results to discrimination of genetic diversity among the basil cultivars and genotypes (figure 2a, b). The constellation plot showed the genetic diversity among the 61 basil genotypes in 2017 (figure 2a). This plot was divided two main groups as A and B. These main groups also divided two sub-groups as A1, A2, B1 and B2. The most of the basil genotypes (49 genotypes) located in group A. The sub-group A1 had the most genotypes (26 genotypes) including Bolu genotype and midnight cultivarcompared to sub-group A2. Twenty-three genotypes located in sub-group A2. The main gropu B included 12 basil genotypes and B1 sub-group had 8 basil genotypes and B2 had only 4 genotypes. B group (B1 and B2 sub-groups) showed differences depending on leaf color than A group (A1 and A2). In experiment 2018 depending UPOV criteria, constellation plot had genetic differences among the 50 basil genotypes in 2018. This plot was also split the two main groups as A and B. These groups were also divided into two main subgroups as A1, A2, and B1 and B2. The first group (group A) had 17 genotypes (two cultivars and 15 basil geniotypes). The subgroup A1 had two cultivars (dino and moonlight) and one genotype and subgroup A2 had 14 genotypes. The second group (group B) had 33 genotypes including two cultivars. The subgroup B1 had two basil cultivars (midnight and large sweet) and 16 genotypes and subgroup B2 had 15 basil genotypes. The constellation plot consisted of 6 clusters and group A had two clusters as C1 and C2, and group B had four clusters as C3, C4, C5 and C6. Basil cultivars took place three different clusters as C1, C3 and C4. Group A2 or C2 had the highest basil genotype counts as well as group A1 or C1 had the lowest basil genotype counts (Figure 2b). 7, 11, 47 basil cultivars and genotype located in group A1 and they showed differences in terms of SH compared the other genotypes. C6 had differences based on PL and FSLI and this cluster took place in different part. It was determined that PH, LBPCS and LBSM properties had no effect to separation the genetic diversity of basil genotypes. These properties were found similar in all basil genotypes.

**Figure 2.**
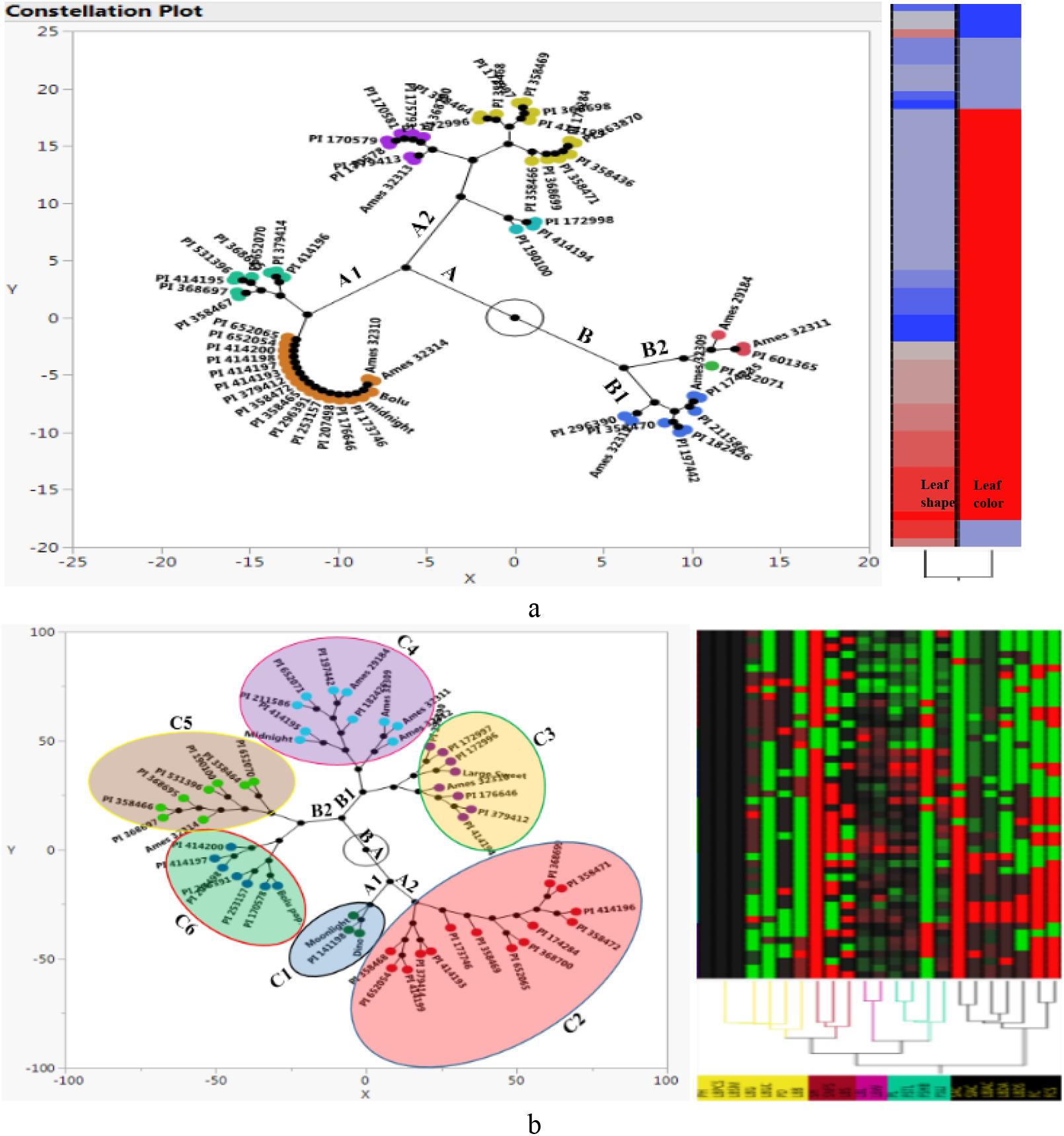
Constellation plot pf basil genotypes (a: 2017 genotypes, b: 2018 genotypes)

Carovic-stanko et al. [16] conducted a study on 27 morphological properties of basil genotypes. It was reported that 46 *Ocimum basilicum* genotypes were divided to 6 clusters depending on 25 morphological properties and same leaf color located different place in dendrogram analysis. Our 2017 and 2018 experimental results showed similar with previous literature.

## 4. Conclusions

On the influence of different genotypes on *Ocimum basilicum* morphology, yield and essential oil values was provided in this study. All morphological parameters evaluated were affected by different *Ocimum basilicum* genotypes. Sizeable variation was observed between individual genotypes which was predominantly due to differences in essential oil and herb yield, and the prospect of selectively breeding for this novel trait. Correlation analysis and PCA showed differences in each year. There was found correlation between yield component and essential oil content in 2017, however in 2018, there was not correlation between these treatments. PCA had totally over than 50% in 2017 and 2018 years. The constellation plot showed genetic diversity and variation appeared depending on leaf shape and color in 2017 and UPOV criteria in 2018. The overall the results of this study was suggested that PI 652070 and PI 296391 genotypes had the highest herbs yield. In addition, PI 358469 and Ames 32309 genotypes had the highest essential oil content. So, genotypes of this later group might be good parents to be used in improvement programs of basil.

## Author Contributions

Conceptualization, writing-review and editing G.Y. supported by M.C.; data curation G.Y. and M.C..; resources G.Y.; methodology G.Y. All authors have read and accepted the published version of the manuscript.

